# High performance of a GPU-accelerated variant calling tool in genome data analysis

**DOI:** 10.1101/2021.12.12.472266

**Authors:** Qian Zhang, Hao Liu, Fengxiao Bu

## Abstract

Rapid advances in next-generation sequencing (NGS) have facilitated ultralarge population and cohort studies that utilized whole-genome sequencing (WGS) to identify DNA variants that may impact gene function. Massive sequencing data require highly efficient bioinformatics tools to complete read alignment and variant calling as the fundamental analysis. Multiple software and hardware acceleration strategies have been developed to boost the analysis speed. This study comprehensively evaluated the germline variant calling of a GPU-based acceleration tool, BaseNumber, using WGS datasets from several sources, including gold-standard samples from the Genome in a Bottle (GIAB) project and the Golden Standard of China Genome (GSCG) project, resequenced GSCG samples, and 100 in-house samples from the China Deafness Genetics Consortium (CDGC) project. Sequencing data were analyzed on the GPU server using BaseNumber, the variant calling outputs of which were compared to the reference VCF or the results generated by the Burrows-Wheeler Aligner (BWA) + Genome Analysis Toolkit (GATK) pipeline on a generic CPU server. BaseNumber demonstrated high precision (99.32%) and recall (99.86%) rates in variant calls compared to the standard reference. The variant calling outputs of the BaseNumber and GATK pipelines were very similar, with a mean F1 of 99.69%. Additionally, BaseNumber took only 23 minutes on average to analyze a 48X WGS sample, which was 215.33 times shorter than the GATK workflow. The GPU-based BaseNumber provides a highly accurate and ultrafast variant calling capability, significantly improving the WGS analysis efficiency and facilitating time-sensitive tests, such as clinical WGS genetic diagnosis, and sheds light on the GPU-based acceleration of other omics data analyses.

## 1 Introduction

Genomic DNA variation represents a critical genetic source of variation that alters protein function and expression, affects human diseases and phenotypes, and can provide markers associated with various populational traits. Next-generation sequencing (NGS) is commonly used to identify DNA variants in a high-throughput manner. Compared to other enrichment-based NGS approaches, whole-genome sequencing (WGS) covers up to 98% of the human genome, providing comprehensive and unbiased variant detection in individuals ^1^. With the development of NGS at a pace far exceeding that predicted by Moore’s Law in recent decades, the cost of sequencing is rapidly decreasing, facilitating the broad application of WGS in clinical and research genetic testing ^2,3^. Ultralarge WGS projects have been initiated, such as Genomics England ^4^, All of Us ^5^, and the China Metabolic Analytics Project ^6^, aiming for in-depth understanding of the underlying molecular mechanisms of human diseases and traits, where accurate identification and interpretation of DNA variants are the foundation. However, due to the enormous amount of sequencing data, the efficiency of variant calling has become a significant bottleneck, with technical complexities remaining.

Conventional variant calling pipelines are often based on CPU servers and open-source software, such as Genome Analysis Toolkit (GATK) ^7^ and VarScan ^8^. GATK HaplotypeCaller calls germline variants through the local de novo assembly of haplotypes in a region showing signs of variation. The pipeline based on GATK Best Practice reportedly completed WGS variant calling on a sample in about 24 hours ^9^. Significant improvements are required to meet the massive needs of WGS studies. A widely used solution is the high-performance cluster (HPC)-based speedup, which reduces the total computation time by using multiple CPU servers to analyze a set of data simultaneously. The use of large clusters implies an increase in the cost of initial equipment procurement, maintenance, and energy consumption. Therefore, it is essential to maximize the utilization of computing resources on a single server to achieve satisfying efficiency. Multiple algorithms and tools for speeding up variant callers have been developed ^10–12^, such as Sentieon DNAseq, a reimplementation of the GATK Best Practice workflow ^13^. By optimizing and recompiling the variant calling algorithms, DNAseq can achieve a 10-fold increase in processing speed while providing results nearly identical to those of the GATK pipeline ^13,14^. However, these CPU computing-based solutions have two major drawbacks: 1) limited parallelism in a CPU environment and 2) the growing gap between CPU computing power and sequencing data throughput. As a promising solution, heterogeneous computing-based approaches have emerged to use different architectures of computing units, such as graphics processing units (GPUs) or field-programmable gate arrays (FPGAs), which are more parallelable than CPUs and more powerful under certain conditions. As an example, Illumina Dragen integrated an FPGA card to boost the variant caller. It also adopted its own bioinformatics algorithms to make good use of the integrated circuits (ICs) on board. The read mapper of Dragen was based on a hash algorithm instead of the Burrows–Wheeler transform (BWT) algorithm used by Burrows-Wheeler Aligner (BWA). In addition, the variant calling algorithm was based on a hidden Markov model, whose running speed can be improved significantly because of its parallelable nature ^15^. However, the concerns of FPGA server-based variant callers include the limited reusability of the dedicated FPGA server for other analytic tasks and hardware-associated upgrading processes. Given their powerful parallel capabilities and wide range of usability, GPU server-based variant calling solutions are gaining more attention ^16–18^. Compared to the GATK pipeline, NVIDIA Clara Parabricks, a GPU-accelerated computational genomics application framework, can reduce the running time of 30X WGS germline analysis on an 8*A100 GPU server by 60-fold ^19^. With a large number of processing units and high memory bandwidth, GPUs can also significantly increase deep learning-based variant callers’ training and inference speed ^20,21^.

The present study aimed to comprehensively evaluate a GPU-based variant caller, BaseNumber (SaileGene Inc, Beijing). Using gold standard samples and an in-house WGS dataset, we compared the BaseNumber to the GATK-based pipeline in terms of efficiency, accuracy, reproducibility, scalability, and energy consumption in germline variant calling on human genome data, providing an overall assessment of this high-throughput variant identification tool.

## 2 Methods

### 2.1 WGS Data Preparation

The WGS data for this study were from four sources: 1) seven standards (HG001 ~ HG007) from the Genome in a Bottle (GIAB) project hosted by the National Institute of Standards and Technology ^22^, 2) four standards (D5, D6, F7, and M8) provided by the Golden Standard of China Genome (GSCG), 3) 24 additional WGS resequenced on the GSCG cell lines in six different DNA sequencing facilities using different sequencing platforms (named the Retested_GSCG samples), and 4) 100 in-house samples from the China Deafness Genetics Consortium (CDGC) project. Reference BAM and VCF files were obtained from GIAB and GSCG. For the in-house data, 100 samples were randomly selected from 1085 subjects of the CDGC project that underwent WGS, representing realistic experimental conditions and outputs. As shown in Table 1 and Table S1, the four GSCG standards had high coverage of 144.2X to 146.5X, while all seven GIAB standards had an ultra-high data volume of 247.1X. The average coverage of the CDGC and Retested_GSCG samples was 48.2X, ranging from 31.0X to 118.0X.

**Table 1.** Summary of WGS samples for the evaluation

| Category | Sample size | Coverage | Evaluation content |
| --- | --- | --- | --- |
| CDGC | 100 | 48.89X | Efficiency, consistency, reproducibility, energy consumption |
| GIAB | 7 | 247.14X | Accuracy, scalability |
| GSCG | 4 | 145.05X | Accuracy, scalability |
| Retested_GSCG | 24 | 45.56X | Accuracy |

### 2.2 Variant Calling Pipelines and Testing Environment

The GATK pipeline followed the widely implemented protocol of best practice that includes Fastp (v0.20.1) for FASTQ preprocessing ^23^, BWA (v0.7.17) for read alignment ^24^, Samtools (v1.9) for BAM sorting ^25^, Picard (v2.23.5) for duplication removal ^26^, and GATK (v4.1.7.0) for base quality score recalibration (BQSR) and variant calling ^27^. The BaseNumber pipeline included Saile-Aligner (SLA, v1.0.3) for FASTQ and BAM processing and Saile-Caller (SLC, v1.0.3) for BQSR and variant calling. Docker (v19.03) was used to pack and implement the pipelines on each server. Genome Reference Consortium Human Build 37 (GRCh37) was used as the reference genome, and corresponding reference files were downloaded from the GATK Resource Bundle (ftp.broadinstitute.org/bundle/).

Two GPU servers and ten high-performance CPU servers were utilized for this study. The detailed configurations of the servers are shown in Table 2. In brief, the GPU_Generic server was configured with 8X NVIDIA Tesla V100 cards and more CPU cores, aiming for common GPU-related computing, such as training of deep learning models. The configuration of the GPU_BaseNumber server was optimized for the BaseNumber pipeline, including improved RAM size and the cooling system. The high-performance CPU (CPU_Generic) servers were designed for generic CPU-related analytic tasks, containing a cost-effectively balanced hardware configuration and no GPU acceleration. UNI-T UT230A-II power sockets were used to monitor the instantaneous power consumption and total power consumption. All testing data were stored on the network-attached storage system, connected with the GPU and CPU servers through the InfiniBand network.

**Table 2.**
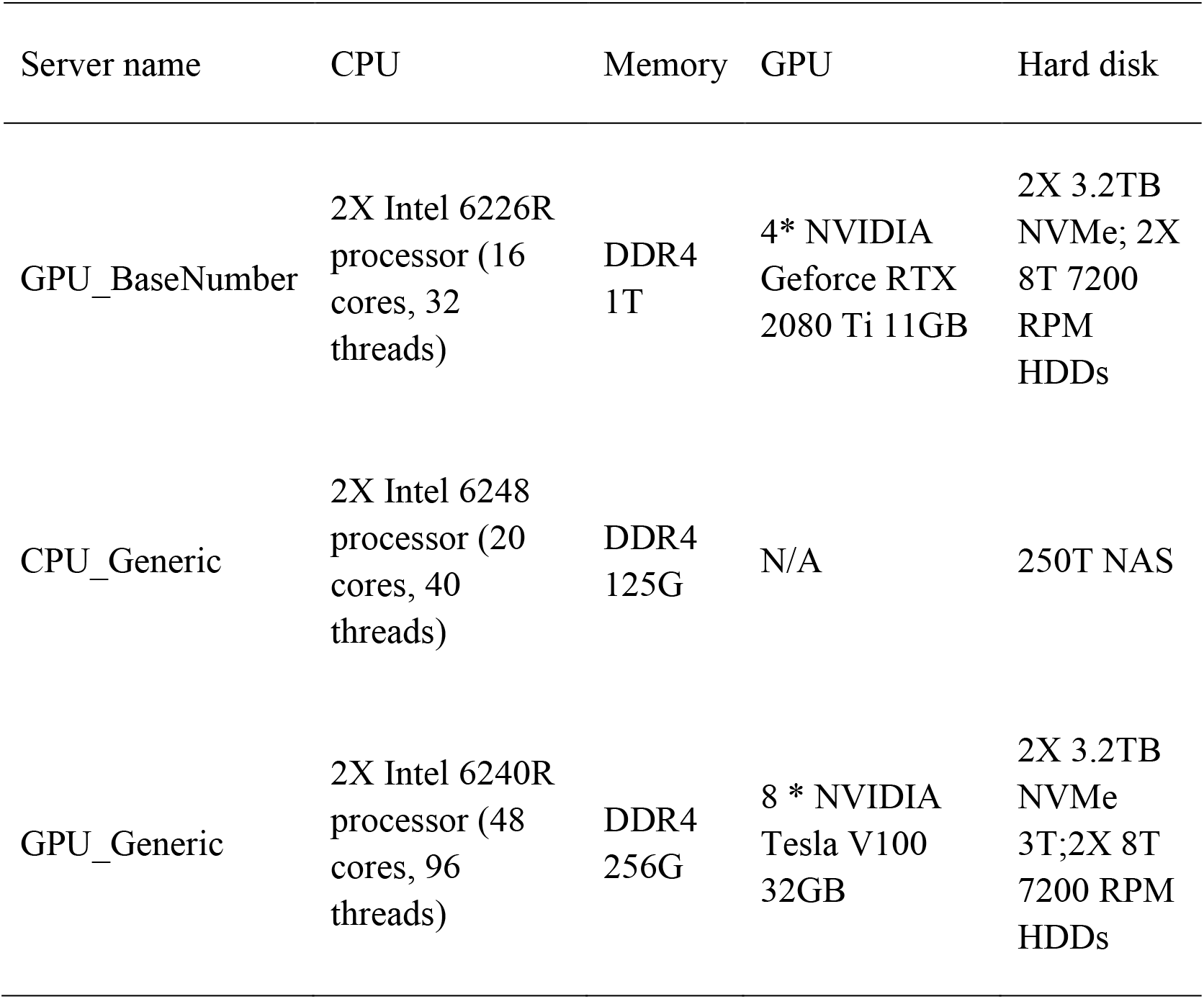
Hardware configuration of testing servers

| Server name | CPU | Memory | GPU | Hard disk |
| --- | --- | --- | --- | --- |
| GPU_BaseNumber | 2X Intel 6226R processor (16 cores, 32 threads) | DDR4 1T | 4* NVIDIA Geforce RTX 2080 Ti 11GB | 2X 3.2TB NVMe; 2X 8T 7200 RPM HDDs |
| CPU_Generic | 2X Intel 6248 processor (20 cores, 40 threads) | DDR4 125G | N/A | 250T NAS |
| GPU_Generic | 2X Intel 6240R processor (48 cores, 96 threads) | DDR4 256G | 8 * NVIDIA Tesla V100 32GB | 2X 3.2TB NVMe 3T;2X 8T 7200 RPM HDDs |

### 2.3 Comprehensive Assessments

The complete evaluation consisted of seven assessments, as illustrated in Figure 1. Gold standard data were applied in the accuracy and scalability assessments, while locally yielded data were used to evaluate the efficiency, consistency, reproducibility, and energy consumption. The accuracy of BaseNumber was assessed using the sequencing data of the Retested_GSCG samples and GIAB standards on the GPU_BaseNumber server. The variant calling output of BaseNumber was compared to the reference VCF files using hap.py from Haplotype Comparison Tools to calculate the precision, recall, and F1 scores of the whole genome and to give genomic regions ^28^. Seqtk was used to randomly select a given proportion of the reads from the original FASTQ files to constitute new FASTQ files with targeted depth ^29^. Raw and downsampled WGS data of GSCG and GIAB standards were used to evaluate the correlation between accuracy and sequencing depth. In addition, downsampled WGS data were applied to assess the correlation between the analysis time and sequencing depth. WGS data of 100 CDGC samples were analyzed on the GPU_BaseNumber and CPU_Generic servers using the BaseNumber and GATK pipelines, respectively, to compare the efficiency. The wall clock time was recorded for each sample and used to calculate the acceleration ratio. BAM and VCF outputs were compared to assess the reproducibility of BaseNumber. Variant calling results of BaseNumber and GATK were compared using HAP for consistency. The analysis times of raw and downsampled HG001 by GPU_BaseNumber and GPU_Generic servers were compared to assess the hardware impact. The GPU_Generic server was configured to assess the influence of the GPU card number on the analysis time.

**Figure 1.**
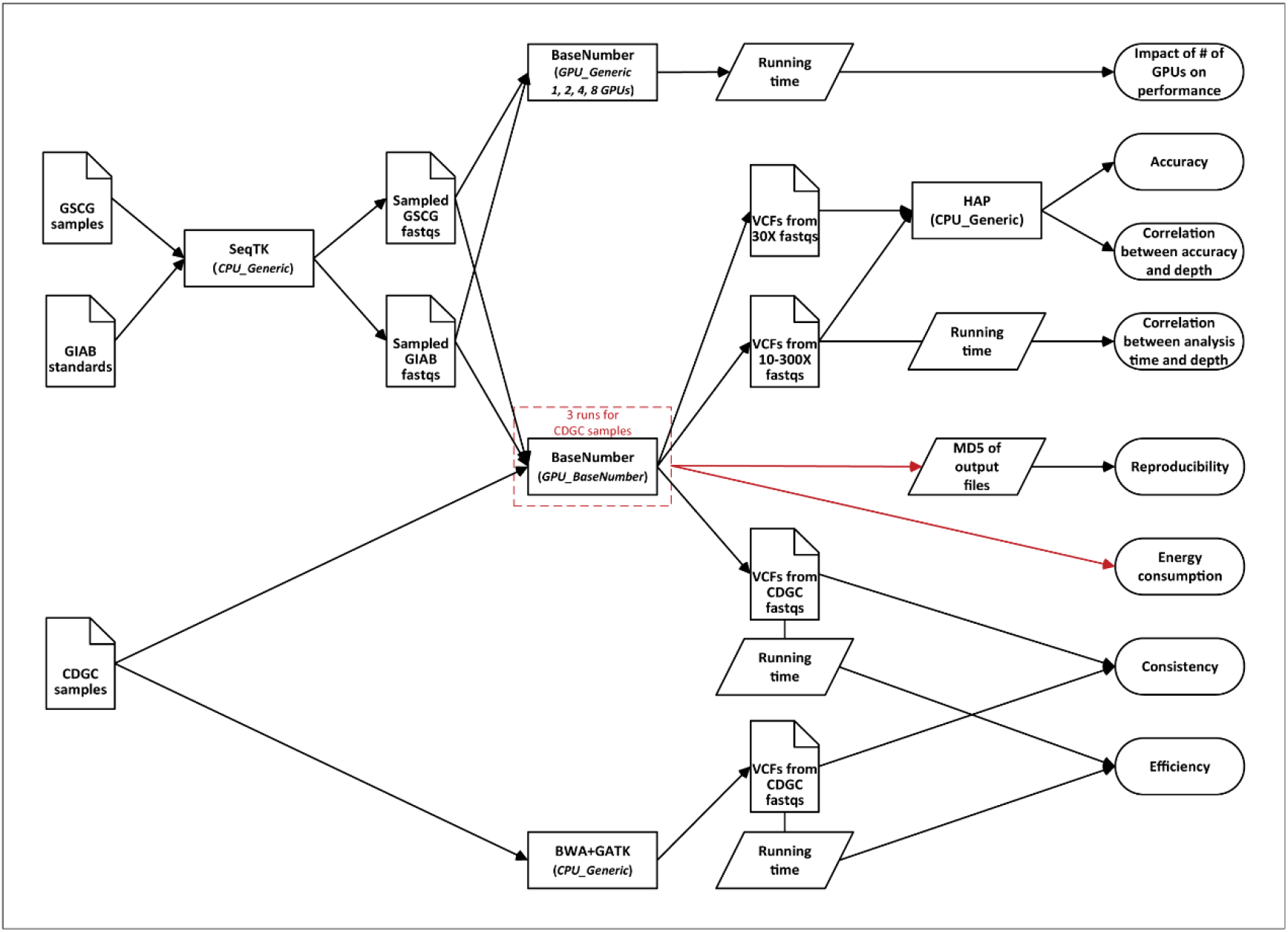
Schema of the study design.

## 3 Results

### 3.1 Accurate Variant Identification using BaseNumber on GSCG and GIAB Standards

We first evaluated the accuracy of the BaseNumber germline variant detection using 24 Retested_GSCG samples and seven GIAB standards (Figure 2A and Table S2). The GIAB standards were downsampled to 30X depth since the raw coverage was excessively high. The reference VCF files of the GSCG and GIAB standards were compared to the BaseNumber variant calling results. For the 31 tested samples, the mean precision rate of all variants was 99.32% (with a standard deviation [SD] of ±0.21%), the mean recall rate was 99.86% (±0.08%), and the mean F1 was 99.59% (±0.10%). Single nucleotide variant (SNV) calling (mean F1 99.63±0.09%) performed slightly better than indel calling (mean F1 99.05±0.09%). We compared the data of the Retested_GSCG samples generated by DNBSEQ-T7 to the same samples generated by NovaSeq 6000. The F1 values were 99.65±0.03% for the DNBSEQ-T7 dataset and 99.56±0.13% for the NovaSeq 6000 dataset, which were comparable. The precision (99.26±0.13%) and recall (99.80±0.09%) rates of the downsampled GIAB samples were also satisfactory.

**Figure 2.**
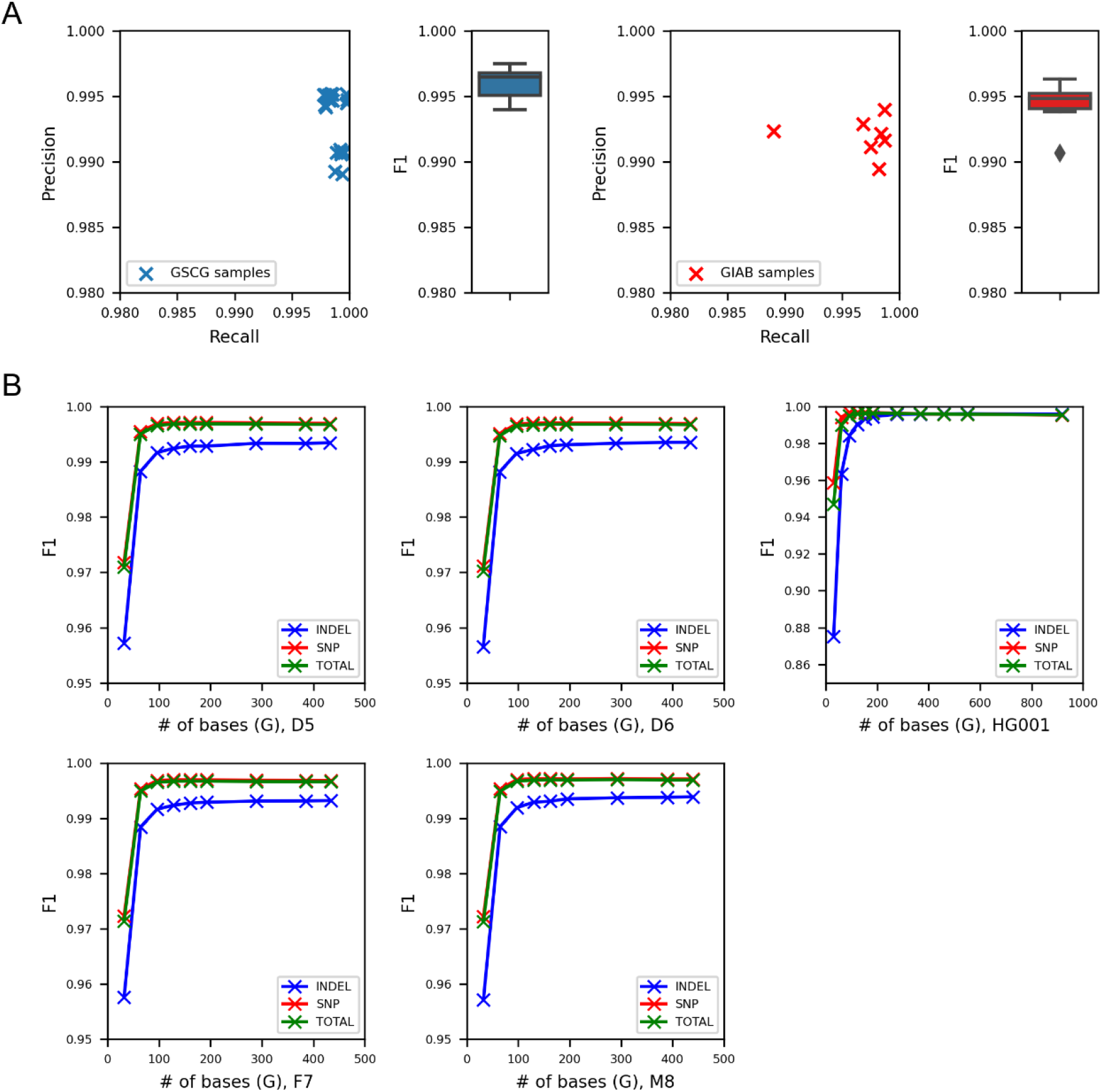
Accuracy evaluation of the BaseNumber using gold standard samples. A) Precision, recall, and F1 score of the BaseNumber on 24 resequenced GSCG and seven GIAB samples. B) F1 score of the BaseNumber rapidly increased along with the sequencing depth.

The correlation between the WGS depth and the variant calling accuracy by BaseNumber was measured with the GSCG standards and GIAB HG001. The reads were randomly retrieved from the FASTQ files with a given proportion to construct a set of simulated WGS samples with gradationally downsampled depth from 10X to 300X (Table S3). The mean F1 was 97.10% for 10X simulated GSCG samples and 94.71% for 10X HG001, but the performance quickly improved with increasing depth (Figure 2B). When the coverage was over 30X, the analysis accuracy reached a stable level even with extremely high coverage of 300X.

### 3.2 Boosted Variant Calling Efficiency of BaseNumber

The wall clock time spent on each step of variant calling on the data of 100 CDGC samples was recorded for the BaseNumber pipeline on the GPU_BaseNumber server and the GATK pipeline on the CPU_Generic servers. As shown in Figure 3A, the mean total analysis time of BaseNumber (from FASTQ to VCF) for each CDGC sample was 23.35±4.75 minutes (ranging from 19.92 to 50.4 minutes). More specifically, the steps of read alignment and polymerase chain reaction (PCR) duplication removal (carried out by SLA) required 10.64±2.68 minutes, the time for generating recalibrated BAM files (carried out by SLC) was 11.59±2.12 minutes, and the variant calling step that outputs gVCF and VCF files (carried out by SLC) required only 1.11±0.28 minutes. As a comparison, a mean of 5018.35±1330.97 minutes (ranging from 3598 to 10179 minutes) was required for GATK to generate final variant calling results from raw sequencing read data of the same sample set. BaseNumber greatly (215.33±37.45 times) improved the computational efficiency of germline variant calling. The analysis time was linearly correlated with the coverage of the WGS data, which ranged from 31X to 118X. The R^2^ of the correlations was 0.863 for BaseNumber and 0.719 for GATK.

**Figure 3.**
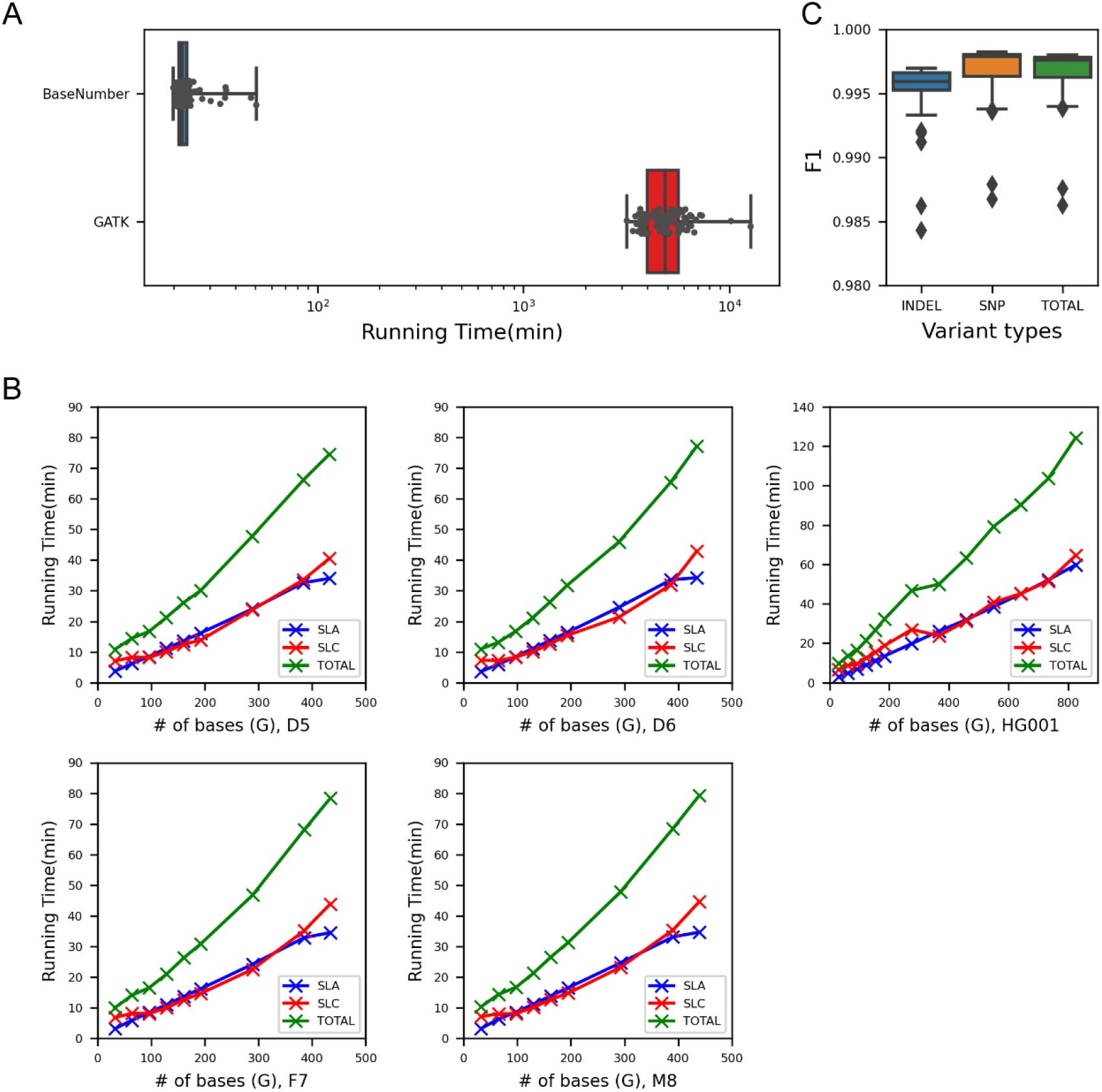
Efficiency of the BaseNumber variant calling process. A) Comparison of analysis time between the BaseNumber and GATK process using 100 CDGC WGS samples. B) Correlation between the BaseNumber analysis time and sequencing depth. C) Variant calling results of the BaseNumber and GATK process were highly consistent

We further explored the effect of data volume on the analysis time using the GSCG and GIAB standards. The FASTQ files of four GSCG standards were randomly downsampled from 10X to 120X. The GIAB HG001 FASTQ was sampled from 10X to 300X. As shown in Figure 3B, a linear correlation between the depth and analysis time was observed even when the coverage was ultrahigh (300X). Interestingly, the R^2^ for SLA was greater than that for SLC, suggesting a stronger linear correlation between the amount of input data and SLA analysis time.

### 3.3 Reproducibility of BaseNumber Outputs

We repeatedly analyzed the CDGC samples to evaluate the reproducibility of BaseNumber. The message-digest algorithm 5 (MD5) values of output files corresponding to the same samples were identical for all three rounds of analysis (Table S4), indicating complete, reproducible, and robust outcomes with BaseNumber. Next, we compared the variant calling results for the CDGC samples from the BaseNumber and GATK pipelines. The precision, recall, and F1 scores were calculated using the corresponding GATK result as the reference for each sample (Figure 3C). The F1 of all variants was on average 99.69±0.19%, ranging from 98.62% to 99.80%. The mean F1 scores were 99.71±0.19% for SNVs and 99.55±0.19% for indels, representing highly similar variant calling outcomes between BaseNumber and GATK on the same sequencing data. We also compared the F1 scores of different sequencing facilities and platforms, and no significant differences were observed.

### 3.4 Impact of the GPU Configuration on the BaseNumber Performance

The configuration of the GPU cards directly impacted the speed of the BaseNumber algorithms. We assessed the running time of the BaseNumber variant caller with different numbers of GPU cards. Downsampled HG001 data (from 10X to 180X) were used for the analysis on the GPU_Generic server configured with eight NVIDIA Tesla V100 cards. As expected, the total analysis time increased as the data volume increased or the number of GPU cards decreased (Figure 4, Table S5). We further dissected the total time into the times for SLA and SLC. The SLA analysis time, which was responsible for alignment and variant calling, continued to decrease as the number of GPU cards increased, while the analysis time of SLC, which included the BQSR process, insignificantly decreased from four GPU cards to eight cards and even increased for HG001_10X. This may be due to IO bottlenecks triggered by SLC when outputting recalibrated BAM files.

**Figure 4.**
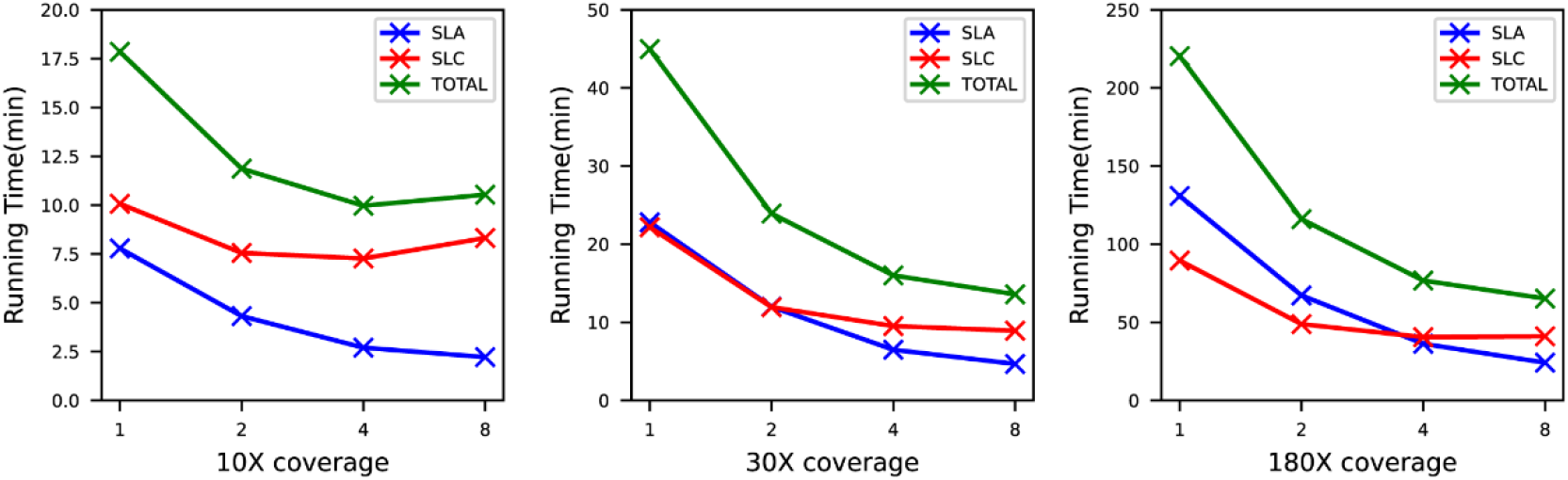
Analysis time of BaseNumber was correlated with GPU configuration and sequencing depth

### 3.5 Energy Consumption

A comparison of the energy consumption of the BaseNumber and GATK pipelines was conducted. Electricity usage was recorded for the variant calling processes of 100 CDGC samples; this was repeated three times. In total, 224 kilowatt-hours were consumed in analyzing these samples three times using BaseNumber on the GPU_BaseNumber server. For each CDGC sample, the energy consumption of BaseNumber was estimated to be 0.746 kilowatt-hours. Similarly, we estimated that the average power consumption per CDGC sample was 33.5 kilowatt-hours for the GATK pipeline on CPU_Generic servers. As a result, the “GPU sever + BaseNumber” solution used 44.9 times less power than the “CPU server + GATK”.

## 4 Discussion

This study comprehensively evaluated the performance of the GPU-accelerated BaseNumber pipeline in germline variant calling. BaseNumber achieved an average F1 value of 99.59% on the gold standards from the GSCG and GIAB projects, in which the accuracy was similar to that of the best-performing variant callers in the PrecisionFDA Consistency Challenge (https://precision.fda.gov/challenges/consistency/results). Additionally, we compared the results of BaseNumber and GATK on CDGC WGS samples with a mean coverage of 48X, and the consistency was remarkable, with an average F1 value of 99.68% using the GATK results as the references. More importantly, we observed an average of 23 minutes taken to analyze the 48X WGS sample using the GPU server, which was more than 200 times faster than the BWA + GATK pipeline. These results show that the BaseNumber pipeline is an attractive alternative to the commonly used BWA + GATK pipeline.

With the emergence of million-sample-sized WGS projects, ultrafast and highly accurate variant calling is essential for further genome analysis. For example, the Genome Aggregation Database (gnomAD) (v3.1) integrated 76,156 WGS samples worldwide, providing invaluable population genetic information to support a wide range of research and clinical applications ^30^. If the average depth of gnomAD samples is 30X, BaseNumber requires only 90 days to complete the variant calling from raw sequencing data using ten servers configured with GPU. Clearly, leveraging the capabilities of BaseNumber can result in significant savings in terms of investment in server hardware, room space, support facilities, and staffing. The reduced analysis time also significantly decreases energy consumption and computing costs. Based on the price of GPU servers and power consumption per sample, we estimate that if 100,000 30X WGS samples are processed in four years, the analysis cost per sample can be less than$0.40. The analysis strategy can be flexibly and finely formulated, facilitated by the significant improvement in efficiency and cost control. Reference genomes, read mapping, variant calling, and quality control algorithms can be tuned at a relatively low sunk cost even as a large cohort study progresses. Moreover, the agile and easy implementation of GPU-based software ensures the operability and timeliness of the pipeline adjustments. During idle time, the GPU server can be used for other high-performance parallel computing tasks, such as deep learning model training, image recognition, and natural language processing, to maximize the usage of the server resources.

The principle of BaseNumber’s germline variant calling algorithm is similar to that of GATK HaplotypeCaller. It reassembles the aligned reads in the active regions where variants may be present to capture SNVs and indels accurately. However, it is challenging to accelerate variant calling on the GPU-based platform. Simple transplantation of HaplotypeCaller to the GPU was reportedly inefficient, with only a 2.3 times speedup ^16,31^. The alignment and variant calling processes were relatively coarse-grained; therefore, the parallelism of corresponding algorithms required reconstruction to support the concurrent operation of thousands of GPU cores. In addition, considering the ultrahigh bandwidth of graphics memory, a dedicated management system needs to be developed to modulate the computing flow. I/O operations have to be optimized to eliminate bottlenecks of the system. With these improvements, BaseNumber achieves high-throughput parallel acceleration and analysis efficiency. Similar performance was claimed for the NVIDIA Clara Parabricks; reportedly, its GPU caller can complete variant analysis of a 30X human WGS in 22 minutes on a DGX A100 server, but Parabricks has not yet revealed detailed information of the evaluation ^19^.

This study focused on the WGS germline variant calling scenario, while BaseNumber is also a proper solution for analyzing ultrahigh coverage sequencing data. Given the high computational capabilities, BaseNumber had a shorter running time, requiring no downsampling processes such as that implemented in GATK to handle excessive coverage regions. As a result, BaseNumber yielded excellent reproducibility in that the outputs for the same data were identical. With the support of the high processing speed of GPUs, some other time-consuming methods, such as graph alignment ^32^ and genotype imputation ^33^, could be applied to boost the accuracy of read mapping and variant calling. BaseNumber might benefit from the adoption of advanced storage systems, such as parallel file systems and flash solid-state drive (SSD) network-attached storage (NAS), and enhanced I/O capabilities to further boost the performance, which became a relative bottleneck in this evaluation.

The results should be interpreted considering several limitations. First, we were unable to compare GATK and BaseNumber on the same GPU server due to the large size of samples for the evaluation and slow process of the GATK pipeline. The GPU_BaseNumber server was configured with better I/O components (such as more RAM and a larger SSD) than the CPU_Generic servers. The performance of GATK might be slightly improved on the GPU_BaseNumber server, but it is unlikely to change the main results. Second, due to the nature of short-read sequencing ^24,34,35^, BaseNumber, like other variant callers based on NGS data, suffered a significant decrease in variant calling accuracy in low mappability regions. Such regions include segmental duplications (segdups) ^36^, tandem repeats ^37^, variable, diversity and joining (VDJ) recombination regions ^38^, and other genomic regions enriched with repetitive sequence. Variant identification and interpretation in these regions depend on the advancement of long-read sequencing techniques.

In conclusion, the GPU-based BaseNumber provides a high accuracy and ultrafast variant calling. It can maximize the efficiency of WGS analysis in large population studies, improve the utilization of hardware resources, and meet the requirements of time-sensitive tests, such as clinical WGS genetic diagnosis. Moreover, BaseNumber shed light on the use of GPUs to accelerate other bioinformatic pipelines based on a similar concept and design ideas to satisfy the increasing demands of multi-omics data analysis, providing powerful support for future individual precision medicine.

## Supporting information

Detailed information of the testing samples

Recall, precision, and F1 scores of BaseNumber in the Retested_GSCG and GIAB samples

Correlation between the accuracy of BaseNumber and sequencing depth

MD5 values of BaseNumber output files from three rounds of analysis

Effects of GPU card counts and sequencing depth on the analysis time

## 6 Author Contributions

F. B., H. L., and Q. Z. conceived the study. H. L. wrote the manuscript, with the contributions by F. B.. Q. Z. organized and performed the evaluation, with contributions by F. B. and H. L.

## 8 Acknowledgments

We are grateful to Prof. Leming Shi provided the GSCG gold standard data.

## Supplementary Appendix

Table S1. Detailed information of the testing samples

Table S2. Recall, precision, and F1 scores of BaseNumber in the Retested_GSCG and GIAB samples.

Table S3. Correlation between the accuracy of BaseNumber and sequencing depth

Table S4. MD5 values of BaseNumber output files from three rounds of analysis

Table S5. Effects of GPU card counts and sequencing depth on the analysis time

